# An atlas of transcription factors expressed in the *Drosophila melanogaster* pupal terminalia

**DOI:** 10.1101/677260

**Authors:** Ben J. Vincent, Gavin R. Rice, Gabriella M. Wong, William J. Glassford, Kayla I. Downs, Jessica L. Shastay, Kenechukwu Charles-Obi, Malini Natarajan, Madelaine Gogol, Julia Zeitlinger, Mark Rebeiz

## Abstract

During development, transcription factors and signaling molecules govern gene regulatory networks to direct the formation of unique morphologies. As changes in gene regulatory networks are often implicated in morphological evolution, mapping transcription factor landscapes is important, especially in tissues that undergo rapid evolutionary change. The terminalia (genital and anal structures) of *Drosophila melanogaster* and its close relatives exhibit dramatic changes in morphology between species. While previous studies have found network components important for patterning the larval genital disc, the networks governing adult structures during pupal development have remained uncharted. Here, we performed RNA-seq in whole *Drosophila melanogaster* terminalia followed by *in situ* hybridization for 100 highly expressed transcription factors during pupal development. We find that the terminalia is highly patterned during pupal stages and that specific transcription factors mark separate structures and substructures. Our results are housed online in a searchable database (flyterminalia.pitt.edu) where they can serve as a resource for the community. This work lays a foundation for future investigations into the gene regulatory networks governing the development and evolution of *Drosophila* terminalia.

**Summary:** We performed RNA-seq in whole *Drosophila melanogaster* terminalia (genitalia and analia) followed by *in situ* hybridization for 100 highly expressed transcription factors during pupal development. We find that the pupal terminalia is highly patterned with specific transcription factors marking separate structures and substructures. Our results are housed online in a searchable database (flyterminalia.pitt.edu) where they can serve as a resource for the community. This work lays a foundation for future investigations into the gene regulatory networks governing the development and evolution of *Drosophila* terminalia.

## Introduction

As animal development proceeds, transcription factors and signaling molecules are expressed in precise patterns to specify cell fate in space and time (Levine and Davidson 2005). These genes ultimately impinge upon cellular effectors, forming gene regulatory networks that alter cellular behavior and generate complex morphologies (Smith *et al.* 2018). Changes within gene regulatory networks can have cellular consequences and result in morphological differences between species (Rebeiz *et al.* 2015). To understand how body parts are built during development and modified through evolution, we must define and dissect their relevant gene regulatory networks.

Of all the anatomical parts in the animal body plan, genitalia have been of particular interest for many evolutionary questions. Genital morphology diverges rapidly between species, which has led some to theorize that males and females are locked in an arms race such that changes in shape or size of genital structures can give one sex a reproductive advantage (Hosken and Stockley 2004; Brennan and Prum 2015) while others theorize that cryptic female choice has led to these morphological differences (Eberhard 1985; Simmons 2014). The accumulation of divergent morphologies between species may then lead to miscoupling of genitalia during interbreeding, reducing viability or fecundity (Masly 2011; Yassin and David 2016; Tanaka *et al.* 2018). Genital morphology is also critical for taxonomic classification, as it is often the only way to reliably distinguish closely related species (Okada 1954; Bock, I.R. & Wheeler, M.R. 1972; Kamimura and Mitsumoto 2011). Previous studies have highlighted several novel genital morphologies that may provide key insights into how new traits evolve (Kopp and True 2002; Yassin and Orgogozo 2013). Despite their intensive study, the molecular basis of genital evolution remains still poorly understood.

The genitalia of *Drosophila melanogaster* and its close relatives provide a unique opportunity to determine how gene regulatory networks build complex and evolving structures. Most previous work on genital development has focused on the larval genital disc, where transcriptomics and targeted genetic experiments have identified several genes that alter adult genitalia when perturbed (Chen and Baker 1997; Gorfinkiel *et al.* 1999; Keisman and Baker 2001; Chatterjee *et al.* 2011). However, much less is known about the genes that control genital development during metamorphosis, when many of the adult structures form through epithelial remodeling (Glassford *et al.* 2015). Quantitative trait locus (QTL) mapping studies have also been performed in *Drosophila*, which have identified several large genomic regions that contribute to genital diversification between crossable sister species (Macdonald and Goldstein 1999; Zeng *et al.* 2000; Masly *et al.* 2011; McNeil *et al.* 2011; Tanaka *et al.* 2015; Takahara and Takahashi 2015). An examination of the gene regulatory networks which govern development of these structures during pupal stages may yield insights into the developmental partitioning of a complex tissue, the causative genes that underlie morphological differences between species, and the origins of novel traits.

The adult male terminalia (comprising both the genitalia and analia) of *D. melanogaster* are subdivided into five main structures, following recently revised nomenclature (Rice *et al.* 2019b): the hypandrium, phallus, surstylus (clasper), epandrial ventral lobe (EVL, also known as the lateral plate), and cercus (also known as the anal plate) (Figure 1A). By 28 hours after puparium formation (APF), four structures can be distinguished in the developing terminalia: hypandrium, phallus, cercus, and the tissue which will give rise to the EVL and surstylus (Figure 1B). By 48 hours APF, the pupal terminalia effectively prefigure adult structures – the surstylus and EVL have separated, and the epandrial posterior lobe has formed along with many other substructures associated with the hypandrium and phallus (Figure 1B). Therefore, in less than 1 day, the pupal terminalia undergo a dramatic remodeling process that builds many adult structures. This rapid transformation motivated our search for transcription factors that pattern these structures during pupal development.

**Figure 1:**
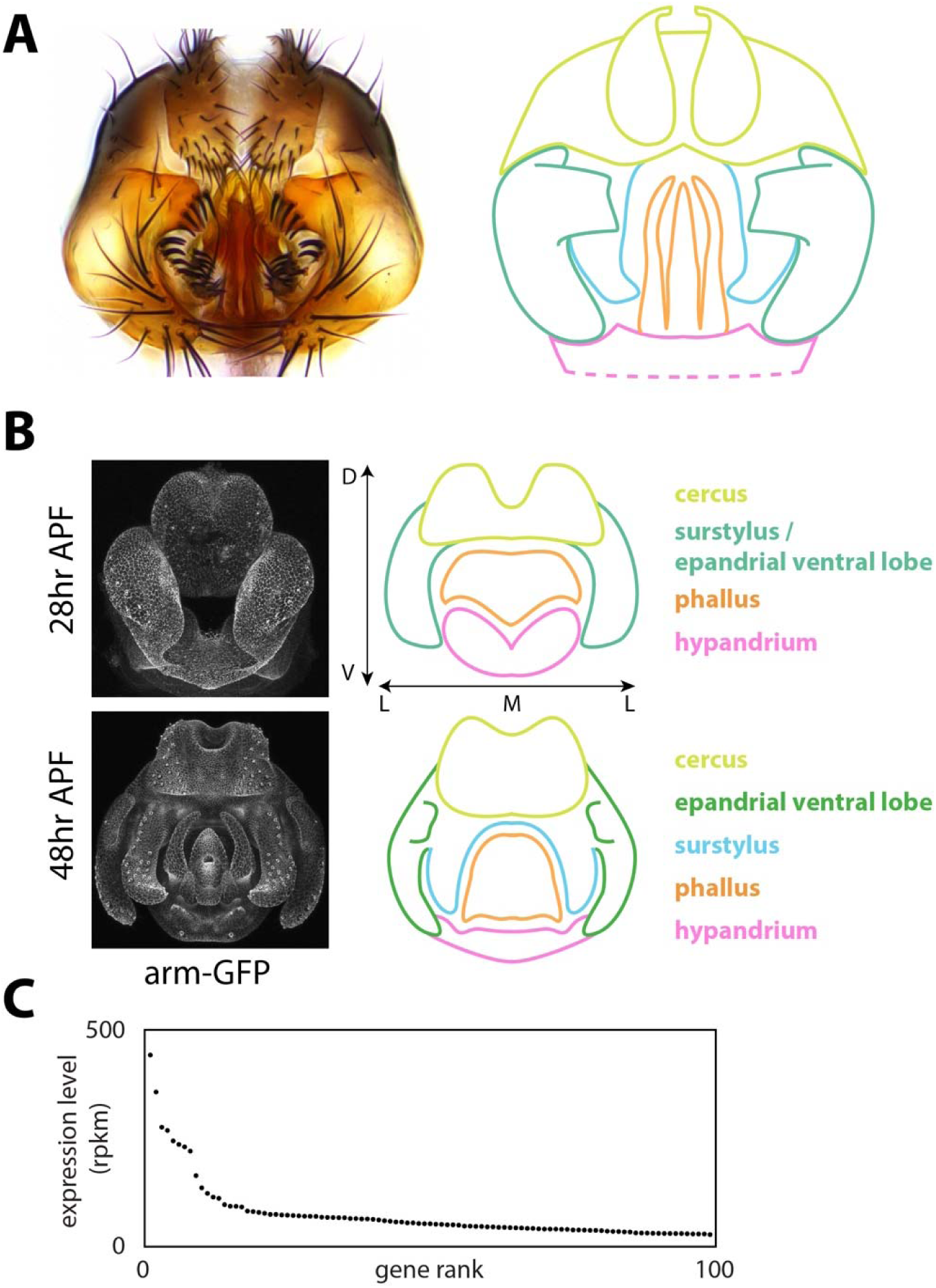
Overview of male terminalia in *Drosophila melanogaster*. A) Left: light microscopy image of adult male terminalia. Right: schematic of major terminal structures. Pink: hypandrium; orange: phallus; green: epandrial ventral lobe; cyan: surstylus; yellow: cercus. The hypandrium extends beyond the cartoon, as represented by dotted lines. Note that our annotations of the cercus includes epandrial dorsal lobe (EDL) and subepandrial sclerite; these are difficult to distinguish during development and thus have been collapsed under the umbrella of cerus structutes. B) Left: confocal microscopy images of developing male terminalia at two developmental time points in a transgenic line where apical cell junctions are fluorescently labeled using an *armadillo-GFP* fusion transgene. Right: schematic of major terminal structures in development, color coded as above. Dorsal-ventral (D-V), and medio-lateral (M-L) axes are labeled. The anterior axis projects into the page while the posterior axis projects out of the page. C) Expression levels of the 100 most highly-expressed transcription factors at 28 hours after puparium formation (APF).

In this study, we performed RNA-seq ofmale terminalia during early pupal development and identified highly expressed transcription factors that may operate during this stage. We then used *in situ* hybridization to build a gene expression atlas of 100 transcription factors in the male pupal terminalia at two time points during development. Most of these genes were highly patterned, especially during the late time point, and we identified genetic markers for many structures and substructures that exhibit morphological differences between Drosophilids. Our data are housed in a searchable online database (flyterminalia.pitt.edu) that will expand as new expression patterns are charted. We believe that the transcription factors characterized here draw the outlines of gene regulatory networks that control genital development and evolution in *Drosophila*.

## Materials and Methods

Detailed, formatted protocols for probe design and synthesis, sample collection, dissection and fixation, and *in situ* hybridization can be found at flyterminalia.pitt.edu.

### RNA-seq and transcriptomic analysis

RNA was isolated from single pupal terminal samples dissected at 24 hours APF or 28 hours APF using the Maxwell® 16 Tissue DNA Purification Kit (Promega). Poly-A RNA-seq libraries were generated using a Clontech library preparation kit (040215). Individual libraries from four different samples were generated for each time point, and libraries were sequenced on an Illumina HiSeq 2500. Sequencing reads from 3 lanes of 51-base Hi-seq data were aligned with tophat (2.0.13) to the dm3 assembly (Trapnell *et al.* 2009), which was retrieved from the UCSC Genome Browser with annotations from Flybase (ftp://ftp.flybase.net/genomes/Drosophila_melanogaster/dmel_r5.57_FB2014_03/gff/). Reads were counted in unioned exons using bedtools count (Quinlan and Hall 2010). Genes expressed in the terminalia were compared to the FlyTF.org list of annotated transcription factors (Pfreundt *et al.* 2010). RNA-seq counts are available at the NCBI Gene Expression Omnibus, accession number GSE133732.

### Probe design and synthesis

Templates for 200-300 basepair RNA probes were designed from a large exon present in all annotated isoforms of each examined gene. Exons were chosen by retrieving the decorated FASTA from flybase.org, and annotated isoforms were examined using the UCSC genome browser. After exon selection, Primer3Plus was used to design PCR primers that would amplify a 200-300 base pair region, and 5-10 candidate primer pairs were screened using the UCSC In Silico PCR tool to identify sets that will amplify the region of interest from the most diverged Drosophilid species possible. This screening process was implemented to maximize the utility of any particular primer set for other species. Reverse primers were designed beginning with a T7 RNA polymerase binding sequence (TAATACGACTCACTATAG), and template DNA was PCR amplified from adult fly genomic DNA extracted using the DNeasy kit (QIAGEN). Digoxigenin-labeled probes were then synthesized using *in vitro* transcription (T7 RNA Polymerase, Promega / Life Technologies), ethanol precipitated, and resuspended in water for Nanodrop analysis. Probes were stored at -20°C in 50% formamide prior to *in situ* hybridization.

### Sample collection, dissection and fixation

Male *D. melanogaster* white pre-pupa (genotype: *yw;*+;+) were collected at room temperature and incubated in a petri dish containing a moistened Kimwipe at 25°C for 28 hours or 48 hours prior to dissection. After incubation, pupae were impaled in their anterior region and immobilized within a glass dissecting well containing cold phosphate buffered saline (PBS). The posterior tip of the pupa (20-40% of pupal length) was separated and washed with a P200 pipette to flush the pupal terminalia into solution. Samples were then collected in PBS with 0.1% Triton-X-100 (PBT) and 4% paraformaldehyde (PFA, E.M.S. Scientific) on ice, and multiple samples were collected in the same tube. Samples were then fixed in PBT + PFA at room temperature for 30 minutes, washed twice in methanol and twice in ethanol at room temperature, and stored at - 20°C.

### In situ hybridization and imaging

We used an InsituPro VSi robot to perform *in situ* hybridization. Briefly, dissected terminalia were rehydrated in PBT, fixed in PBT with 4% PFA and prehybridized in hybridization buffer for 1 hr at 65°C. Samples were then incubated with probe for 16h at 65°C before washing with hybridization buffer and PBT. Samples were blocked in PBT with 1% bovine serum albumin (PBT+BSA) for 2 hours. Samples were then incubated with anti-digoxigenin Fab fragments conjugated to alkaline phosphatase (Roche) diluted 1:6000 in PBT+BSA. After additional washes, color reactions were performed by incubating samples with NBT and BCIP (Promega) until purple stain could be detected under a dissecting microscope. Samples were mounted in glycerol on microscope slides coated with poly-L-lysine and imaged at 20X or 40X magnification on a Leica DM 2000 with a Leica DFC540 C camera. For most images available online, extended focus compilations were acquired using the ImageBuilder module of the Leica Application Suite.

In interpreting our results, we performed several qualitative comparisons to increase our confidence in the data. First, we processed samples from both time points simultaneously in the same basket and staining well. For many genes, we observed uniform expression in 28h samples but patterned expression in 48h samples. These observations gave us confidence that the uniform early expression was not due to background staining. Similarly, we occasionally observed expression patterns in samples from one time point but not the other, which fostered confidence that the absence of expression was not due to experimental failure. As an additional safeguard, we compared results from different genes stained in the same batch to detect cross-contamination. Finally, we compared equivalent samples in annotating our results, such that the representative images presented in this manuscript were corroborated by replicates.

## Results

### Global measurements of gene expression levels in early pupal terminalia

To identify transcription factors that may play a role in genital and anal development, we performed RNA-seq on early pupal terminalia dissected at 24 hours and 28 hours after puparium formation (APF). We found that 11,816 genes are expressed at levels greater than 1 read per kilobase per million reads (rpkm) in at least 1 time point, including 282 annotated transcription factors (Pfreundt *et al.* 2010). Among the 100 most highly expressed transcription factors at 28 hours APF, the expression levels ranged from 442 to 27 rpkm (Figure 1C). These genes formed the basis for our gene expression atlas.

### An atlas of the genital transcription factor landscape

Our transcriptomic analysis suggested that a large number of transcription factors are expressed in the pupal terminalia. In order to glean spatial and temporal expression information for these candidates, we performed *in situ* hybridization (ISH) in pupal terminalia at 28 hours and 48 hours APF. ISH measurements are qualitative and variable – distinguishing signal from background can be challenging, especially for genes that are uniformly expressed, and results may vary between biological replicates. We addressed these challenges through several comparisons (see Materials and Methods). In addition to the results presented here, our full dataset is housed online at flyterminalia.pitt.edu. We built this database to increase the accessibility, transparency and reproducibility of our results. We include full protocols for our methods as well as key experimental details underlying the results for each experiment. For each gene, we also include annotations of all tissues in which evidence of gene expression was observed. Finally, to accurately represent the variability in our results, this database includes images of all samples that met the quality control standards of our experimental pipeline.

For the remainder of the manuscript, we organize our results by describing select transcription factors expressed in each structure of the terminalia.

### The Epandrial Ventral Lobe (Lateral Plate)

The epandrial ventral lobe (EVL, also called the lateral plate) is a periphallic structure lateral to the phallus (Rice *et al.* 2019b). The epandrial posterior lobe (hereafter referred to as the posterior lobe) develops from the EVL (Glassford *et al.* 2015) and is a key diagnostic feature of the *melanogaster* clade (Coyne 1983; Markow and O’Grady 2005). Multiple groups have attempted to map the genomic regions associated with morphological changes in the posterior lobe (Macdonald and Goldstein 1999; Zeng *et al.* 2000; Masly *et al.* 2011; McNeil *et al.* 2011; Tanaka *et al.* 2015; Takahara and Takahashi 2015). In addition, a previous study identified a gene regulatory network associated with posterior lobe development that also functions in the development of the posterior spiracle, a larval structure involved in gas exchange (Glassford *et al.* 2015). Multiple transcription factors within the posterior lobe network appeared among our candidates, and we used these genes as positive controls for our methods.

At 28h APF, the tissue that will form the surstylus and the EVL exists as a single continuous epithelium that later undergoes cleavage to form both structures by 48h APF (Figure 1B, (Glassford *et al.* 2015). Hereafter, we refer to this single structure as the epandrial ventral lobe / surstylus (EVL/S). In accordance with previous results, we found that *Pox neuro* (*Poxn*) is expressed in the EVL/S at 28h APF and the EVL at 48h APF (Figure 2A). In addition to *Poxn*, we found that *Abdominal-B* and *empty spiracles* are expressed in the EVL/S and EVL, as well as within the posterior lobe domain (Figure 2C-E); both genes were previously identified as posterior lobe network components (Glassford *et al.* 2015).

**Figure 2:**
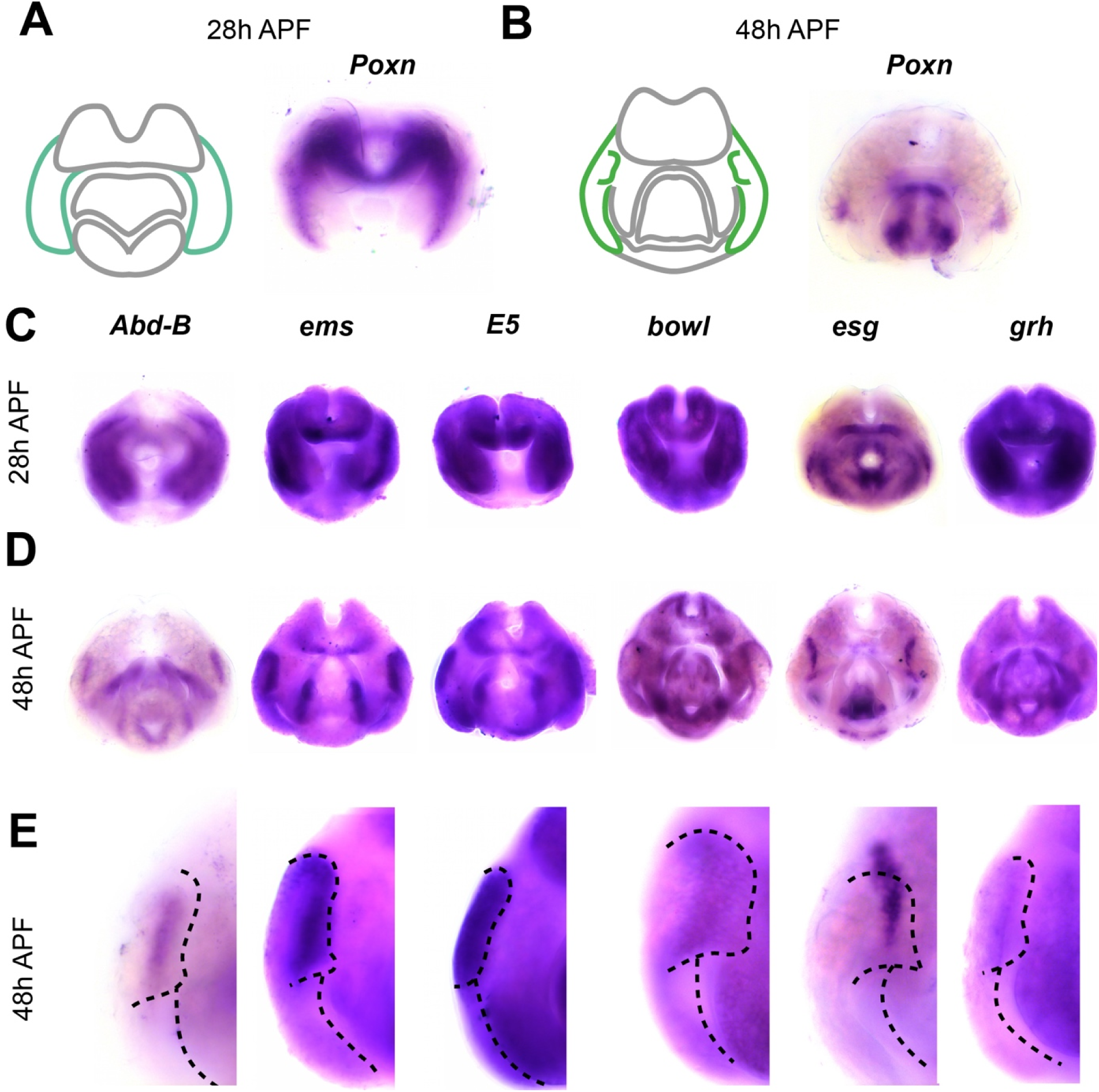
Transcription factors expressed in the epandrial ventral lobe (EVL). A) Left: schematic of major terminal structures at 28 hours APF with the epandrial ventral lobe / surstylus highlighted in turquoise. Right: Light microscopy image of *in situ* hybridization data for *Pox neuro* mRNA at 28 hours APF. Purple signal indicates localization of target mRNA. B) Left: schematic of major terminal structures at 48 hours APF with the EVL and posterior lobe highlighted in green. Right: Light microscopy image of *in situ* hybridization data for *Poxn* mRNA at 48 hours APF. Additional *In situ* hybridization data for EVL-specific factors at 28 hours APF (C) 48 hours (D), and in closeups at 48h (E). The boundaries of the posterior lobe and the medial boundary of the EVL are indicated by dashed lines.

In addition to these known factors, we identified many other transcription factors expressed in the EVL and posterior lobe. We found that *E5* is expressed in the posterior lobe, the ventral portion of the EVL (see additional samples online), and the phallus. *E5* is a homeodomain transcription factor (Dalton *et al.* 1989) associated with variation in posterior lobe morphology among *Drosophila melanogaster* populations (Takahashi *et al.* 2018). We also found that *brother of odd with entrails limited* (*bowl*) is expressed in the posterior lobe at 48 hours APF, as well as other tissues throughout the terminalia (Figure 2C-E). *bowl* is a target of Notch signalling and has been previously implicated in leg development and epithelial rearrangements in the hindgut (Iwaki *et al.* 2001; de Celis Ibeas and Bray 2003).

In addition to genes localized within the posterior lobe, we found that *escargot* (*esg*) and *grainyhead* (*grh*) are expressed in the EVL at both timepoints, but occupy a compartment medial to the posterior lobe – both are expressed near the location where EVL tissue separate from the surstylus (Figure 2C-E). *esg* is a *snail*-related transcription factor that functions in the development of larval imaginal discs (Whiteley *et al.* 1992; Hayashi *et al.* 1993; Fuse *et al.* 1996), while *grh* is associated with the maternal-zygotic transition during embryonic development, as well as morphogenetic processes in several developmental contexts (Hemphälä *et al.* 2003; Narasimha *et al.* 2008; Harrison *et al.* 2010).

We did not identify a transcription factor that serves as a unique/non-ambiguous marker for the EVL or the posterior lobe – all genes expressed in the EVL were also expressed in at least one other tissue (Figure 2C and D). For example, *Abd-B, ems, E5* and *esg* accumulate mRNA in the posterior lobe and phallus, but within different phallic substructures (Figure 2C, see below for descriptions of phallic morphology). *grh* and *bowl* are also expressed in other, distinct terminal structures (Figure 2C). Thus, transcription factors expressed in these structures are not unique, but show patterns of co-expression which differ from factor to factor.

### The Surstylus (Clasper)

The surstylus (also known as the clasper) is a curled outgrowth located medial to the EVL (Rice *et al.* 2019b). Like the posterior lobe, the surstylus exhibits morphological differences between closely related species in the *melanogaster* subgroup (Bock, I.R. & Wheeler, M.R. 1972), and has been the focus of quantitative trait locus (QTL) mapping efforts (True *et al.* 1997; Tanaka *et al.* 2015). A recent study identified *tartan*, a cell adhesion protein, as a gene that contributes to changes in surstylus morphology between *Drosophila simulans* and *Drosophila mauritiana* (Hagen *et al.* 2018). However, while RNAi experiments in *Drosophila melanogaster* have identified several genes that influence surstylus morphology (Tanaka *et al.* 2015), little is known about the gene regulatory network that governs its development during pupal stages.

We found that *odd-paired* (*opa*) is expressed exclusively in the surstylus at 48h APF, as well as the medial portion of the EVL/S at 28h APF (Figure 3A and B). These data suggest that *opa* is a surstylus-specific marker, and can also identify presumptive surstylus tissue prior to its cleavage from the EVL. In other tissues, *opa* controls the formation of parasegment boundaries during embryogenesis (Clark and Akam 2016), as well as morphogenetic events in the formation of the midgut and head (Cimbora and Sakonju 1995; Lee *et al.* 2007).

**Figure 3:**
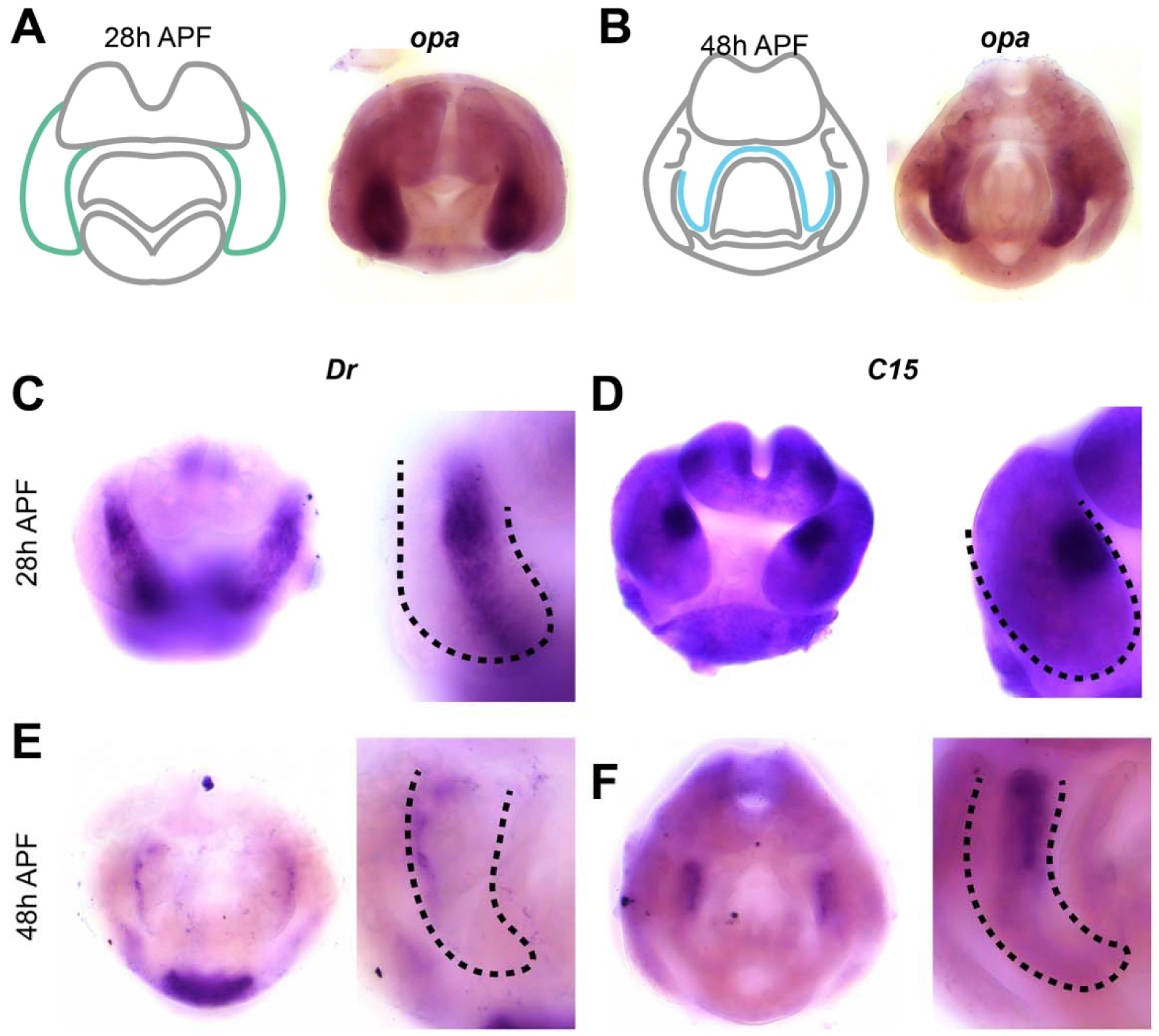
Transcription factors expressed in the surstylus. A) Left: schematic of major terminal structures at 28 hours APF with the epandrial ventral lobe / surstylus indicated in turquoise. Right: Light microscopy image of *in situ* hybridization data for *odd-paired* mRNA at 28 hours APF. B) Left: schematic of major terminal structures at 48 hours APF with the surstylus outlined in cyan. Right: Light microscope image of *in situ* hybridization data for *odd-paired* mRNA at 48 hours APF. (C-D) *in situ* hybridization data for *Drop* mRNA in whole terminalia (left) and at higher magnification (right) at 28 hours APF (C) and at 48 hours APF (D). (E-F) *in situ* hybridization data for *C15* mRNA in whole terminalia (left) and at higher magnification (right) at 28 hours APF (E) and 48 hours APF (F). Dashed lines indicate the boundary of the EVL/surstylus (C and D) or the surstylus (E and F).

In addition to *opa*, we found transcription factors expressed in specific subcompartments of the surstylus. *Drop* (*Dr*) is expressed in presumptive surstylus tissue at 28h APF, as well as a more restricted compartment at 48h APF, which may represent the boundary between the surstylus and the EVL (Figure 3, C and E). *Dr* has been previously implicated in genital development and is expressed in larval (L3) genital discs (Chatterjee *et al.* 2011). We also found that *C15* is expressed in a dorsal-medial compartment of the presumptive surstylus at 28h APF, as well as at the base of the surstylus at 48h APF (Figure 3, D and F). *C15* functions in the development of the amnioserosa during embryogenesis (Rafiqi *et al.* 2008), as well as during leg development where it interacts with *apterous* and *bowl* (Campbell 2005), both of which exhibit patterned expression in the pupal terminalia (see flyterminalia.pitt.edu). These data show that like the EVL, the surstylus can be delineated into subcompartments by the expression patterns of transcription factors during pupal development.

### The Cercus (Anal plate)

The cercus (anal plate) is composed of two flat, semicircular sheets of cuticle on the dorsal side of the terminalia (Rice *et al.* 2019b). The cercus is derived from abdominal segment 10 while the rest of the male terminalia originates from abdominal segment 9 (Keisman *et al.* 2001). This structure shows dramatic variation in bristle number and morphology within and between Drosophilid species (Lachaise *et al.* 1981; Kopp and True 2002), which in some cases have been implicated in reproductive incompatibility (Tanaka *et al.* 2018). QTL analysis for differences in the total cercus area between *D. mauritiana* and *D. simulans* identified causative genomic regions, but were unable to resolve these to the level of individual genes (True *et al.* 1997; Tanaka *et al.* 2015). We note that our annotations of genes expressed in the cercus may include expression patterns that localize to the developing epandrial dorsal lobe (EDL) and subepandrial sclerite. In the pupal terminalia, the cercus, subepandrial sclerite, and EDL are continuously joined, and their boundaries are unclear, however when possible we differentiate them below.

We found that *caudal* (*cad*) was expressed throughout the cercus at both time points, as well as the tissue that connects the surstyli together (subepandrial sclerite) at 48h APF (Figure 4, A and B). We did not observe *cad* expression in other structures; thus *caudal* serves as a marker for these tissues at this stage of development. *cad*, which functions in the anterior-posterior patterning network in embryogenesis (Macdonald and Struhl 1986; Rivera-Pomar *et al.* 1995; Olesnicky *et al.* 2006; Vincent *et al.* 2018), has been previously implicated in the development of the cercus and interacts with the genes *Distaless* (*Dll*) and *brachyenteron* (*byn*) in the L3 genital disc (Moreno and Morata 1999).

**Figure 4:**
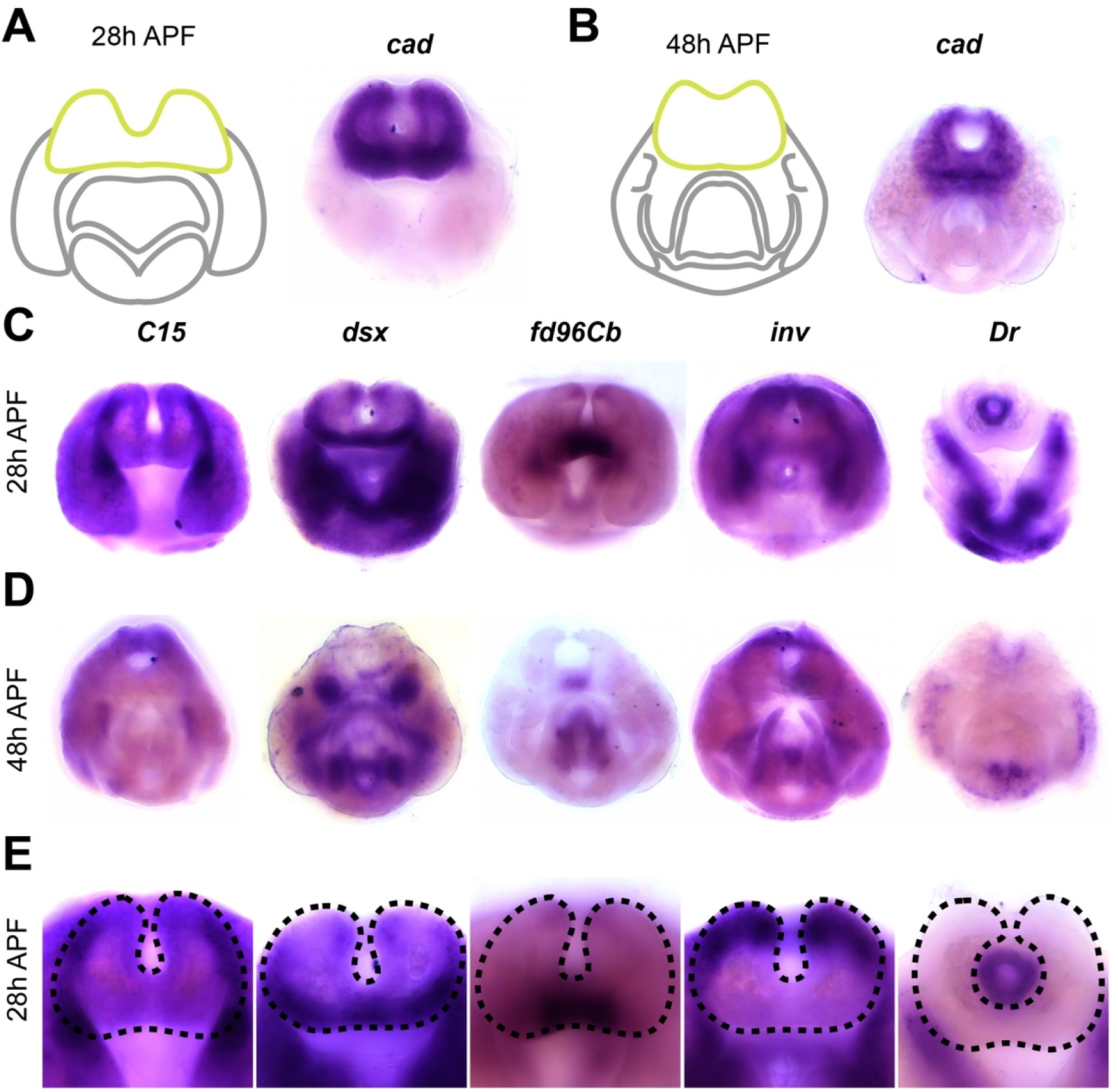
Transcription factors expressed in the cercus. A) Left: schematic of major terminal structures at 28 hours APF with the cercus indicated in yellow. Right: Light microscopy image of *in situ* hybridization data for *cad* mRNA at 28 hours APF. B) Left: schematic of major terminal structures at 48 hours APF with the cercus indicated in yellow. Right: Light microscopy image of *in situ* hybridization data for *cad* mRNA at 48 hours APF. *In situ* hybridization data for transcription factors *fd96Cb, C15, dsx, inv, Dr* at 28 hours APF (C), 48h (D), and in closeups at 28h (E). Dashed lines indicate the boundary of the cercus.

We also identified several transcription factors that are expressed in distinct subcompartments of the cercus. *C15* was expressed in the lateral boundaries, while *doublesex* (*dsx*) was expressed on the anterior-ventral face. *dsx* is a known regulator of sexually dimorphic traits (Hildreth 1965; Baker and Ridge 1980). *forkhead domain 96Cb* (*fd96Cb)* was expressed only in the medial portion of the ventral side in a pattern that clearly resolves by 48h APF. *invected* (*inv*) was expressed on the dorsal and lateral sides along with *engrailed*; these genes are partially redundant in other tissues and specify the anterior compartment of other abdominal segments (Kopp *et al.* 1997), including the terminalia (Epper and Sánchez 1983; Chen and Baker 1997; Casares *et al.* 1997). Finally, several genes are expressed in the developing rectum, including *Dr* (Figure 3C-E), *knirps*, and *tramtrack* (see flyterminalia.pitt.edu).

### The Hypandrium

The hypandrium is a plate-like structure that flanks the phallus on the ventral side (Rice *et al.* 2019b). The hypandrium contains several substructures, including the hypandrial phragma, medial gonocoxite, pregonites, lateral gonocoxites, and (Figure 1C). Within the hypandrium, the lateral gonocoxite and the pregonites exhibit rapid evolution across Drosophilids (Okada 1954; Kamimura and Mitsumoto 2011). While few genes have been previously implicated in hypandrial development, genetic perturbations in *Dr* cause changes in hypandrial morphology (Chatterjee *et al.* 2011), and one study localized the loss of hypandrial bristles to a *cis*-regualtory element of the *scute* gene (Nagy *et al.* 2018).

We found that *Dichaete* (*D*) is expressed in the hypandrial phragma (i.e. deep into the sample when viewed from the posterior) at both time points (Figure 5A and B). *D* is a member of the *Sox* family of transcription factor genes and is critical in embryogenesis (Russell *et al.* 1996). We also found that several transcription factors are expressed in hypandrial substructures. For example, *Dr* is expressed throughout the medial gonocoxite and weakly in the hypandrial phragma (Figure 5D). In contrast, *esg* is localized to the base of the pregonites as well as the posterior tip of the lateral gonocoxite (Figure 5E). Taken together, we found discrete gene expression patterns within the pupal domains of or the annotated hypandrial substructures.

**Figure 5.**
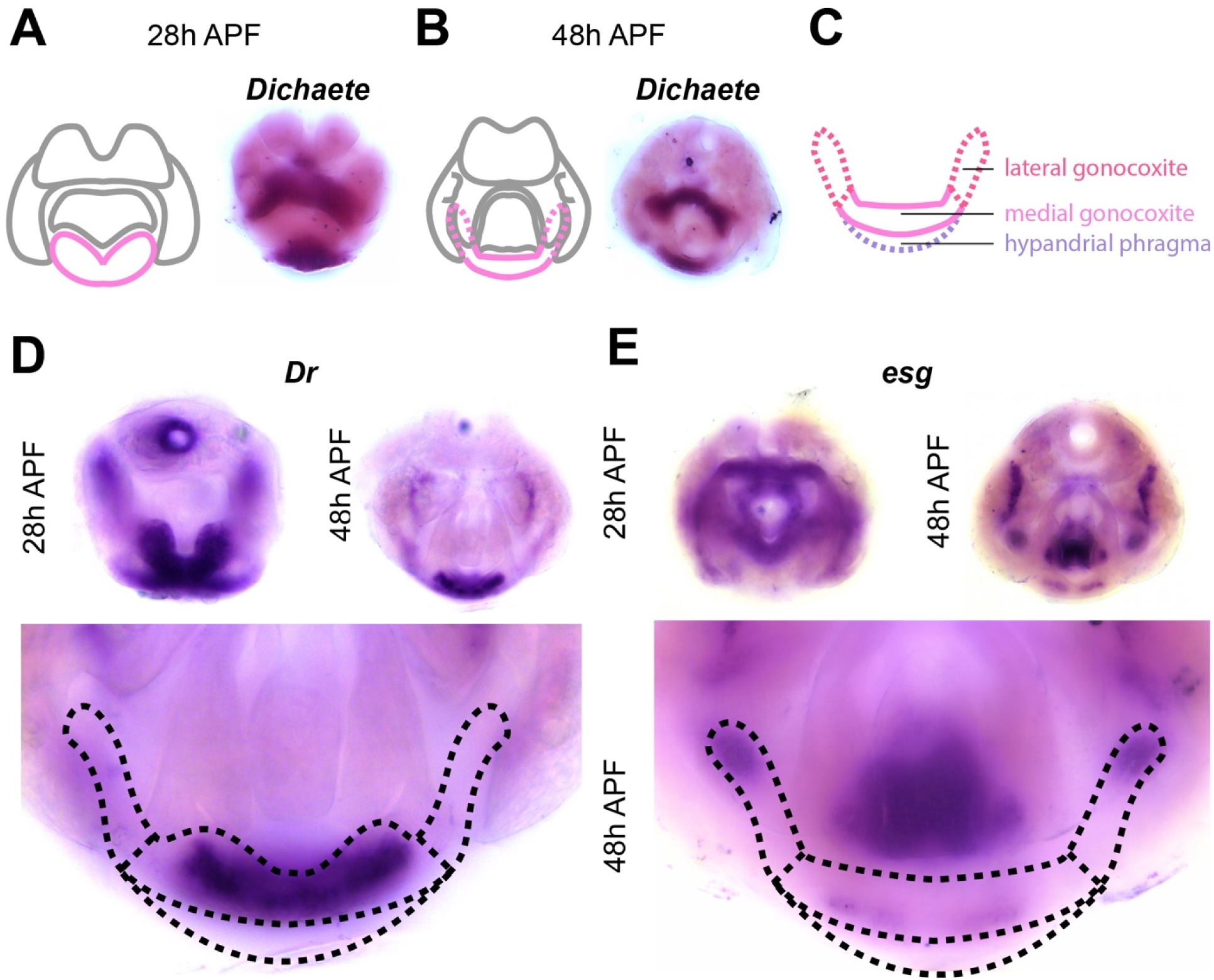
Transcription factors expressed in the hypandrium. A) Left: schematic of major terminal structures at 28 hours APF with the hypandrium indicated in pink. Right: Light microscopy image of *in situ* hybridization data for *Dichaete* (*D*) mRNA at 28 hours APF. B) Left: schematic of major terminal structures at 48 hours APF with the hypandrium indicated in pink. Right: Light microscopy image of *in situ* hybridization data for *D* mRNA at 48 hours APF. C) Cartoon representation of the substructures of the hypandrium: hypandrial phragma (purple), medial gonocoxite (pink), and lateral gonocoxite (red). Dashed lines indicate substructures that are obscured by other parts of the terminalia. *in situ* hybridization data at Left: 28 hours APF, Right: 48 hours APF (right), and Bottom: high magnification images of 48hr APF samples to illustrate details of hypandrial expression patterns for *Dr* (D) and *esg* (E). The boundaries of substructures are indicated by dashed lines.

### The Phallus

The phallus is the male genital organ used for intromission and is composed of four substructures: aedeagus, aedaegal sheath, dorsal postgonites, and ventral postgonites (Rice *et al.* 2019b). Each of these substructures exhibits morphological changes within the *melanogaster* species group (Okada 1954; Kamimura and Mitsumoto 2011), and QTL mapping has identified genomic regions associated with some of these differences (Peluffo *et al.* 2015). Here, we confirmed that *Poxn* is expressed throughout the phallus (Figure 6D), which is consistent with previous observations that *Poxn* is essential for phallic development (Boll and Noll 2002; Glassford *et al.* 2015).

**Figure 6:**
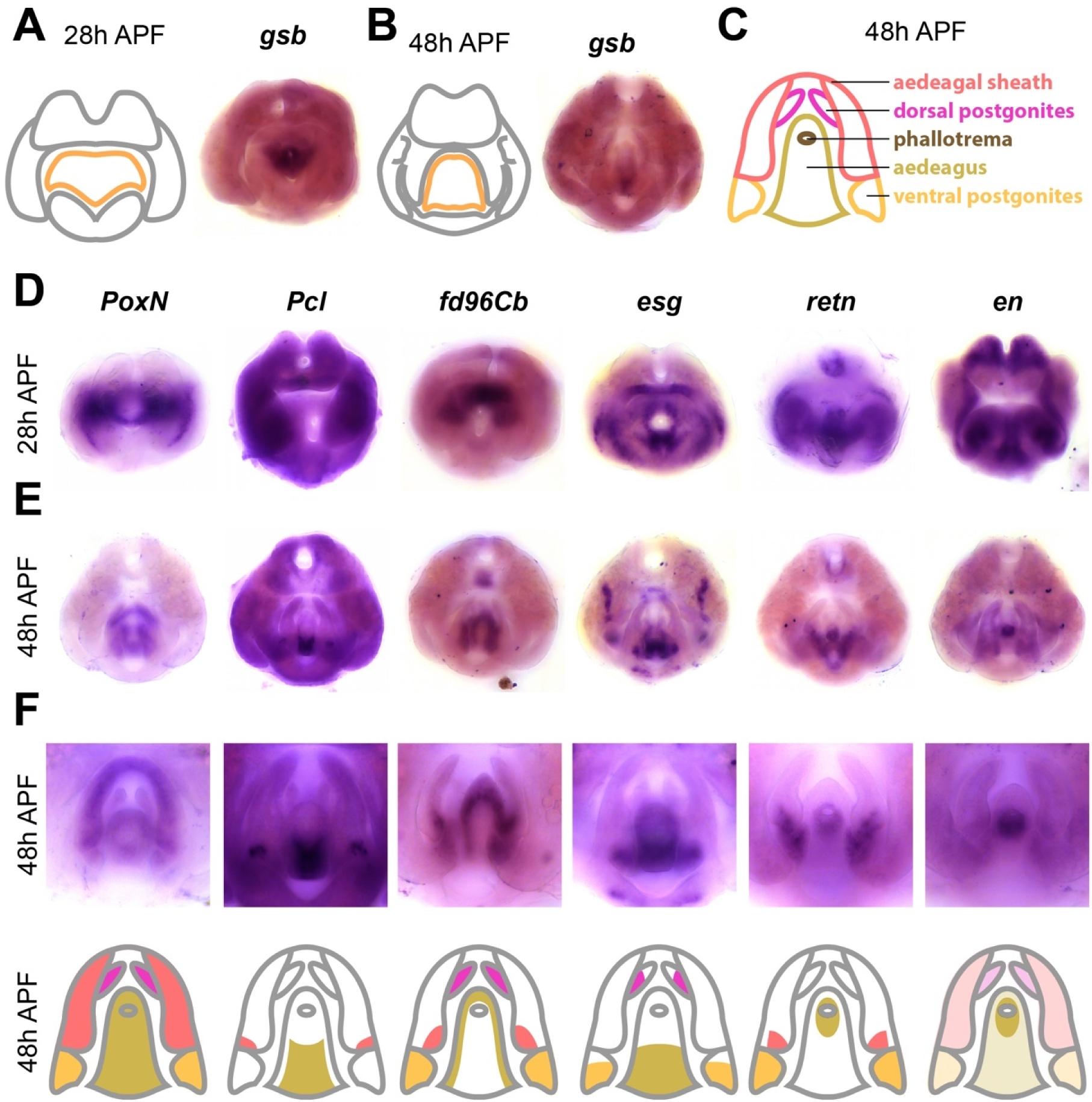
Transcription factors expressed in the phallus. A) Left: schematic of major terminal structures at 28 hours APF with the phallus indicated in orange. Right: Light microscope image of *in situ* hybridization data for *gsb* mRNA at 28 hours APF. B) Left: schematic of major terminal structures at 48 hours APF with the phallus indicated in orange. Right: Light microscopy image of *in situ* hybridization data for *gsb* mRNA at 48 hours APF. C) Cartoon representation of the substructures of the phallus: ventral postgonite (orange), aedeagus (yellow), phallotrema (brown), dorsal postgonites (pink), and aedeagal sheath (red). Additional *In situ* hybridization data for transcription factors *PoxN, esg, fd96Cb, retn, Plc*, and *en* at 28 hours APF (D) and 48 hours APF (E). F) Top: High magnification images of the samples shown in (E) to illustrate details of phallus expression patterns. Bottom: Cartoon representation of the substructures of the phallus, with shading indicating expression within each substructure. Note that for *en*, light shading indicates weak expression throughout the phallus.

The aedeagus is a phallic structure that delivers sperm and exhibits a needle-like shape in *D. melanogaster*. We identified genes that are expressed along the dorsal-ventral axis of the aedeagus in what appear to be non-overlapping patterns. We found that *gooseberry* (*gsb*) was exclusively expressed in the ventral portion of the aedeagus at both 28 and 48hrs APF (Figure 6A and B). *gsb* was previously found to be expressed in the anterior-ventral edge in L3 genital discs (Freeland and Kuhn 1996), and is a segment polarity gene that interacts with *wingless* during embryogenesis (Li and Noll 1993). We also found that *Polycomb-like* (*Pcl*) was expressed in the same compartment as *gsb* at 48h APF, but exhibits broader expression at 28h APF (Figure 6D–F). Reciprocally, we found that *fd96Cb* was expressed in the dorsal portion of the aedeagus. Finally, we identified genes expressed in other aedeagal subcompartments. For example, we found that *esg* was restricted to the anterior base of the aedeagus, while *retained* (*retn*), *inv* and *en* are expressed in the opening of the aedeagus, known as the phallotrema.

The aedeagal sheath along with the dorsal and ventral postgonites are two phallic substructures situated lateral to the aedeagus (Figure 6C). The aedeagal sheath consists of two flat, shield-like extensions that bilaterally flank the aedeagus. We found that several genes were expressed in the sheath, including *fd96Cb* and *retn*. The dorsal and ventral postgonites are two pairs of spike-like extensions that project from the aedeagal sheath. We found that *esg* is expressed at the base of both pairs of postgonites, while *fd96Cb* was expressed throughout the entire structure of both pairs of postgonites. We also found that *retn* (Figure 6F) and *dsx* (flyterminalia.pitt.edu) are expressed in the ventral postgonites, but not the dorsal pair, and we note that *dsx* has a known enhancer that drives expression in this region (Rice *et al.* 2019a). Taken together, we identified genes that are expressed in distinct phallic structures, as well as within subcompartments of individual structures.

## Discussion

In this study, we profiled the transcriptome of the male pupal terminalia in *D. melanogaster* at critical timepoints when major adult structures form. We then determined the spatiotemporal gene expression patterns of the 100 most highly expressed transcription factors during this stage. We identified transcription factors that were expressed in the five major terminal structures, as well as several substructures that exhibit morphological diversity between species. We discuss the implications of our results for the development and evolution of terminalia in Drosophilids.

### Drosophila terminalia as a model system

To appreciate the transformative power of a gene expression atlas, we need to look no further than the *Drosophila melanogaster* embryo. Beginning with the iconic Heidelberg screen (Nüsslein-Volhard and Wieschaus 1980; Wieschaus and Nüsslein-Volhard 2016), which identified genes that control embryonic patterning, many groups have contributed to the development and dissemination of genetic resources for studies in embryogenesis. These resources include transcriptomic profiling (Lott *et al.* 2011) and expression atlases of nearly all genes detectable during this stage of development (Tomancak *et al.* 2007; Lécuyer *et al.* 2007). Quantitative gene expression atlases are now available at cellular resolution for multiple genetic backgrounds and in different species (Fowlkes *et al.* 2008, 2011; Pisarev *et al.* 2009; Staller *et al.* 2015; Karaiskos *et al.* 2017). These atlases enable computational models of gene regulatory networks and enhancer function that have provided insights into the evolution of patterning networks (Wunderlich *et al.* 2012; Wotton *et al.* 2015). However, these resources have revealed that the gene regulatory network which patterns the embryo evolves slowly, producing subtle quantitative changes in gene expression even between distantly related Drosophilids (Fowlkes *et al.* 2011; Wunderlich *et al.* 2019). In contrast, the terminalia contain multiple rapidly evolving structures which can illuminate important and under-explored aspects of gene regulatory network evolution.

We envision this atlas of 100 transcription factors as a first step towards building a comprehensive system for the study of developmental network function and evolution. Our RNA-seq data suggest that additional transcription factors are expressed at 28 hours APF, and it is possible that transcriptomic measurements at other time points or with different methods will reveal additional candidates. We will continue to add additional gene expression measurements to FlyTerminalia (flyterminalia.pitt.edu) as these candidates are pursued. In particular, our atlas provides a foundation for performing and analyzing single-cell RNA-seq experiments on developing pupal terminalia. While single-cell RNA-seq data provide more highly-resolved information on cell types, they do not contain anatomical information on the spatial organization of those cell types. We therefore anticipate that this atlas will permit one to annotate and interpret single-cell RNA-seq data. In the future, we hope to expand FlyGenitalia to include expression patterns in the developing female terminalia, which are historically understudied (Hosken and Stockley 2004; Ah-King *et al.* 2014), as well as expression measurements in other species. By continuing to develop these resources, we hope that *Drosophila* terminalia will become a premiere model system to address many questions in evolution and development.

### Implications for genital evolution

Most of the recent work on the genetic basis of genital evolution has been confined to variation within species and between crossable species (True *et al.* 1997; Zeng *et al.* 2000; Masly *et al.* 2011; Tanaka *et al.* 2015, 2018; Peluffo *et al.* 2015). However, even for the most extensively studied genital traits, only a portion of the heritable changes have been resolved to the level of individual genes (Hagen *et al.* 2018; Nagy *et al.* 2018). This atlas may thus provide useful candidates for numerous unresolved QTL peaks. In addition, many traits evolve on macroevolutionary time scales, excluding the possibility of QTL analysis. Previous work used a comparative analysis of gene expression to identify a network of genes that was co-opted to the posterior lobe – a novel trait restricted to the *melanogaster* clade (Glassford *et al.* 2015). However, the *D. melanogaster* clade contains other unique traits, including structures whose gene regulatory networks have not been previously characterized. In this study, we found several genes that are expressed in lateral gonocoxite (*esg, inv, en*), and postgonites (*esg, fd96Cb, crp, mod, retn* and *dsx*), both of which exhibit morphological changes between species. Furthermore, a ventral postgonite enhancer was recently identified for the gene *doublesex* (Rice et al. 2019a) which may be a useful gene expression driver to manipulate this structure in the future. Other enhancers that drive expression in the larval genital disc may persist in the pupal terminalia and may serve as drivers to target other structures (Jory *et al.* 2012). To assess the functional roles of individual genital structures in copulation, genetic disruption may help complement other techniques such as laser ablation (Polak and Rashed 2010; Kamimura and Polak 2011; LeVasseur-Viens *et al.* 2015).

Rapid morphological changes between species hamper the identification of homology in the genitalia and cercus. Structural homology has previously been defined by similarities in adult morphology, but structures that appear similar may nevertheless not be related by common descent. As a result, there are conflicting claims of homology – the same structure in one species has been called homologous to different structures in other species (Frank et; Grimaldi 1987; Grimaldi David A 1990). Based on our results, we suggest that gene expression profiles may be useful in reconciling conflicting claims of homology. For example, homology is difficult to establish for the postgonites, often referred to as parameres or branches (Kamimura 2007; Yassin and Orgogozo 2013; Peluffo *et al.* 2015). Here, we identified genes that are expressed in both pairs of postgonites (*fd96Cb*, and *esg*), which may help to define homologous structures in other species.

### Implications for genital development

In mapping the transcription factor landscape in the pupal terminalia, we have begun defining the gene regulatory networks that operate in the development of these structures. Identifying relevant transcription factors and measuring their gene expression patterns is an important first step, but we must also determine how these genes interact. At this point, we can infer regulatory interactions by looking for incidences of co-expression or reciprocal expression. For example, it would be interesting to test whether transcription factors expressed in the entirety of particular structures, such as the surstylus marker *odd-paired*, are required for expression of other genes deployed in more restricted subcompartments, such as *C15*. Some of these genes have known regulatory interactions in other contexts, such as *apterous, C15*, and *bowl* (Campbell 2005). While this atlas can be a tool for generating hypotheses about how these gene regulatory networks are wired, these hypotheses must ultimately be rigorously tested via genetic perturbation.

Locating the regulatory DNA that controls these expression patterns will also be critical for defining relevant gene regulatory networks. One notable feature of our results is that most of the identified transcription factors are expressed in multiple locations throughout the pupal terminalia, especially at 48h APF. It remains unclear whether these patterns are controlled by multiple regulatory elements, or if disparate patterns are generated by the same enhancer region (Small *et al.* 1996). It is possible that the enhancers controlling these patterns also operate in other tissues or at different developmental stages (Noon *et al.* 2018; Sabarís *et al.* 2019), as is the case for the posterior lobe enhancer of *Pox neuro* (Glassford *et al.* 2015) and the hypandrial enhancer of *scute* (Nagy et al. 2018). By finding the regulatory sequences that control these gene expression patterns, we can determine the direct targets of transcription factors in this system.

Epithelial remodeling is a critical component of many developmental events, including gastrulation, neural tube formation, and organogenesis (Neumann and Affolter 2006). Studying these processes in *Drosophila* tissues, such as the wing disc and the trachea, has yielded insights into similar processes in mammals (Affolter *et al.* 2003). We focus here on patterned transcription factors because morphogenetic processes are tightly regulated at the level of gene expression. However, we are ultimately interested in the connections between transcription factors and the effectors that ultimately dictate cell behavior (Smith *et al.* 2018). Recent work has implicated a variety of cellular mechanisms in the formation of genital structures, including changes in cell size and cell intercalations in the developing ovipositor (Green *et al.* 2019) and the influence of the apical extracellular matrix in the developing posterior lobe (Smith, et al. submitted). In the future, we hope to characterize the functional roles of transcription factors in both cellular dynamics and adult morphology, and elucidate how the expression and function of these genes are tuned to generate new or different structures over evolutionary time.

## Acknowledgments

We thank the entire Rebeiz lab for assistance optimizing the *in situ* hybridization protocol and helpful discussions, especially Eden McQueen and Donya Shodja for comments on the manuscript. Artoo Situ deserves special recognition for autonomous assistance with *in situ* hybridization. We also thank the University of Pittsburgh Center for Research Computing, especially Kim Wong for technical help on building and hosting website. This project was supported by the Stowers Institute for Medical Research and the National Institutes of Health (GM107387 to MR, 1F32GM130034 to BJV and 1F32GM125329 to WJG).

## References

Affolter M., S. Bellusci, N. Itoh, B. Shilo, J.-P. Thiery, et al., 2003 Tube or not tube: remodeling epithelial tissues by branching morphogenesis. Dev. Cell 4: 11–18.

Ah-King M., A. B. Barron, and M. E. Herberstein, 2014 Genital evolution: why are females still understudied? PLoS Biol. 12: e1001851.

Baker B. S., and K. A. Ridge, 1980 Sex and the single cell. I. On the action of major loci affecting sex determination in Drosophila melanogaster. Genetics 94: 383–423.

Bock, I.R. & Wheeler, M.R., 1972 The Drosophila melanogaster species group. University of Texas Publication 7213: 1–102.

Boll W., and M. Noll, 2002 The Drosophila Pox neuro gene: control of male courtship behavior and fertility as revealed by a complete dissection of all enhancers. Development 129: 5667–5681.

Brennan P. L. R., and R. O. Prum, 2015 Mechanisms and Evidence of Genital Coevolution: The Roles of Natural Selection, Mate Choice, and Sexual Conflict. Cold Spring Harb. Perspect. Biol. 7: a017749.

Campbell G., 2005 Regulation of gene expression in the distal region of the Drosophila leg by the Hox11 homolog, C15. Dev. Biol. 278: 607–618.

Casares F., L. Sánchez, I. Guerrero, and E. Sánchez-Herrero, 1997 The genital disc of Drosophila melanogaster. : I. Segmental and compartmental organization. Dev. Genes Evol. 207: 216–228.

Celis Ibeas J. M. de, and S. J. Bray, 2003 Bowl is required downstream of Notch for elaboration of distal limb patterning. Development 130: 5943–5952.

Chatterjee S. S., L. D. Uppendahl, M. A. Chowdhury, P.-L. Ip, and M. L. Siegal, 2011 The female-specific doublesex isoform regulates pleiotropic transcription factors to pattern genital development in Drosophila. Development 138: 1099–1109.

Chen E. H., and B. S. Baker, 1997 Compartmental organization of the Drosophila genital imaginal discs. Development 124: 205–218.

Cimbora D. M., and S. Sakonju, 1995 Drosophila midgut morphogenesis requires the function of the segmentation gene odd-paired. Dev. Biol. 169: 580–595.

Clark E., and M. Akam, 2016 Odd-paired controls frequency doubling in Drosophila segmentation by altering the pair-rule gene regulatory network. Elife 5. https://doi.org/10.7554/eLife.18215

Coyne J. A., 1983 Genetic basis of differences in genital morphology among three sister species of Drosophila. Evolution 37: 1101–1118.

Dalton D., R. Chadwick, and W. McGinnis, 1989 Expression and embryonic function of empty spiracles: a Drosophila homeo box gene with two patterning functions on the anterior-posterior axis of the embryo. Genes Dev. 3: 1940–1956.

Eberhard W. G., 1985 Sexual Selection and Animal Genitalia Epper F., and L. Sánchez, 1983 Effects of engrailed in the genital disc of Drosophila melanogaster. Dev. Biol. 100: 387–398.

Fowlkes C. C., C. L. L. Hendriks, S. V. E. Keränen, G. H. Weber, O. Rübel, et al., 2008 A quantitative spatiotemporal atlas of gene expression in the Drosophila blastoderm. Cell 133: 364–374.

Fowlkes C. C., K. B. Eckenrode, M. D. Bragdon, M. Meyer, Z. Wunderlich, et al., 2011 A conserved developmental patterning network produces quantitatively different output in multiple species of Drosophila, (A. Kopp, Ed>.). PLoS Genet. 7: e1002346.

Frank et M., Manual of nearctic Diptera.

Freeland D. E., and D. T. Kuhn, 1996 Expression patterns of developmental genes reveal segment and parasegment organization of D. melanogaster genital discs. Mech. Dev. 56: 61–72.

Fuse N., S. Hirose, and S. Hayashi, 1996 Determination of wing cell fate by the escargot and snail genes in Drosophila. Development 122: 1059–1067.

Glassford W. J., W. C. Johnson, N. R. Dall, S. J. Smith, Y. Liu, et al., 2015 Co-option of an Ancestral Hox-Regulated Network Underlies a Recently Evolved Morphological Novelty. Dev. Cell 34: 520–531.

Gorfinkiel N., L. Sánchez, and I. Guerrero, 1999 Drosophila terminalia as an appendage-like structure. Mech. Dev. 86: 113–123.

Green J. E., M. Cavey, E. Médina Caturegli, B. Aigouy, N. Gompel, et al., 2019 Evolution of Ovipositor Length in Drosophila suzukii Is Driven by Enhanced Cell Size Expansion and Anisotropic Tissue Reorganization. Curr. Biol. https://doi.org/10.1016/j.cub.2019.05.020

Grimaldi D. A., 1987 Phylogenetics and taxonomy of Zygothrica (Diptera: Drosophilidae). Filogenia y taxonomía de Zygothrica (Diptera: Drosophilidae). Bull. Am. Mus. Nat. Hist. 186: 104–268.

Grimaldi David A, 1990 A phylogenetic, revised classification of genera in the Drosophilidae (Diptera). Bull. Am. Mus. Nat. Hist. 1–139.

Hagen J. F. D., C. C. Mendes, K. M. Tanaka, P. Gaspar, M. Kittelmann, et al., 2018 tartan underlies the evolution of male genital morphology. bioRxiv 462259.

Harrison M. M., M. R. Botchan, and T. W. Cline, 2010 Grainyhead and Zelda compete for binding to the promoters of the earliest-expressed Drosophila genes. Dev. Biol. 345: 248–255.

Hayashi S., S. Hirose, T. Metcalfe, and A. D. Shirras, 1993 Control of imaginal cell development by the escargot gene of Drosophila. Development 118: 105–115.

Hemphälä J., A. Uv, R. Cantera, S. Bray, and C. Samakovlis, 2003 Grainy head controls apical membrane growth and tube elongation in response to Branchless/FGF signalling. Development 130: 249–258.

Hildreth P. E., 1965 Doublesex, recessive gene that transforms both males and females of Drosophila into intersexes. Genetics 51: 659–678.

Hosken D. J., and P. Stockley, 2004 Sexual selection and genital evolution. Trends Ecol. Evol. 19: 87–93.

Iwaki D. D., K. A. Johansen, J. B. Singer, and J. A. Lengyel, 2001 drumstick, bowl, and lines are required for patterning and cell rearrangement in the Drosophila embryonic hindgut. Dev. Biol. 240: 611–626.

Jory A., C. Estella, M. W. Giorgianni, M. Slattery, T. R. Laverty, et al., 2012 A survey of 6,300 genomic fragments for cis-regulatory activity in the imaginal discs of Drosophila melanogaster. Cell Rep. 2: 1014–1024.

Kamimura Y., 2007 Twin intromittent organs of Drosophila for traumatic insemination. Biol. Lett. 3: 401–404.

Kamimura Y., and H. Mitsumoto, 2011 Comparative copulation anatomy of the Drosophila melanogaster species complex (Diptera: Drosophilidae). Entomol. Sci. 14: 399–410.

Kamimura Y., and M. Polak, 2011 Does surgical manipulation of Drosophila intromittent organs affect insemination success? Proc. Biol. Sci. 278: 815–816.

Karaiskos N., P. Wahle, J. Alles, A. Boltengagen, S. Ayoub, et al., 2017 The Drosophila embryo at single-cell transcriptome resolution. Science 358: 194–199.

Keisman E. L., and B. S. Baker, 2001 The Drosophila sex determination hierarchy modulates wingless and decapentaplegic signaling to deploy dachshund sex-specifically in the genital imaginal disc. Development 128: 1643–1656.

Keisman E. L., A. E. Christiansen, and B. S. Baker, 2001 The sex determination gene doublesex regulates the A/P organizer to direct sex-specific patterns of growth in the Drosophila genital imaginal disc. Dev. Cell 1: 215–225.

Kopp A., M. A. Muskavitch, and I. Duncan, 1997 The roles of hedgehog and engrailed in patterning adult abdominal segments of Drosophila. Development 124: 3703–3714.

Kopp A., and J. R. True, 2002 Evolution of male sexual characters in the oriental Drosophila melanogaster species group. Evol. Dev. 4: 278–291.

Lachaise D., F. Lemeunier, and M. Veuille, 1981 Clinal Variations in Male Genitalia in Drosophila teissieri Tsacas. Am. Nat. 117: 600–608.

Lécuyer E., H. Yoshida, N. Parthasarathy, C. Alm, T. Babak, et al., 2007 Global analysis of mRNA localization reveals a prominent role in organizing cellular architecture and function. Cell 131: 174–187.

Lee H., B. G. Stultz, and D. A. Hursh, 2007 The Zic family member, odd-paired, regulates the Drosophila BMP, decapentaplegic, during adult head development. Development 134: 1301–1310.

LeVasseur-Viens H., M. Polak, and A. J. Moehring, 2015 No evidence for external genital morphology affecting cryptic female choice and reproductive isolation in Drosophila. Evolution 69: 1797–1807.

Levine M., and E. H. Davidson, 2005 Gene regulatory networks for development. Proc. Natl. Acad. Sci. U. S. A. 102: 4936–4942.

Li X., and M. Noll, 1993 Role of the gooseberry gene in Drosophila embryos: maintenance of wingless expression by a wingless--gooseberry autoregulatory loop. EMBO J. 12: 4499–4509.

Lott S. E., J. E. Villalta, G. P. Schroth, S. Luo, L. A. Tonkin, et al., 2011 Noncanonical compensation of zygotic X transcription in early Drosophila melanogaster development revealed through single-embryo RNA-seq, (R. S. Hawley, Ed.). PLoS Biol. 9: e1000590.

Macdonald P. M., and G. Struhl, 1986 A molecular gradient in early Drosophila embryos and its role in specifying the body pattern. Nature 324: 537–545.

Macdonald S. J., and D. B. Goldstein, 1999 A quantitative genetic analysis of male sexual traits distinguishing the sibling species Drosophila simulans and D. sechellia. Genetics 153: 1683–1699.

Markow T. A., and P. O’Grady, 2005 Drosophila: A Guide to Species Identification and Use. Academic Press.

Masly J. P., J. E. Dalton, S. Srivastava, L. Chen, and M. N. Arbeitman, 2011 The genetic basis of rapidly evolving male genital morphology in Drosophila. Genetics 189: 357–374.

Masly J. P., 2011 170 Years of “Lock-and-Key”: Genital Morphology and Reproductive Isolation. Int. J. Evol. Biol. 2012. https://doi.org/10.1155/2012/247352

McNeil C. L., C. L. Bain, and S. J. Macdonald, 2011 Multiple Quantitative Trait Loci Influence the Shape of a Male-Specific Genital Structure in Drosophila melanogaster. G3 1: 343–351.

Moreno E., and G. Morata, 1999 Caudal is the Hox gene that specifies the most posterior Drosophile segment. Nature 400: 873–877.

Nagy O., I. Nuez, R. Savisaar, A. E. Peluffo, A. Yassin, et al., 2018 Correlated Evolution of Two Copulatory Organs via a Single cis-Regulatory Nucleotide Change. Curr. Biol. 28: 3450–3457.e13.

Narasimha M., A. Uv, A. Krejci, N. H. Brown, and S. J. Bray, 2008 Grainy head promotes expression of septate junction proteins and influences epithelial morphogenesis. J. Cell Sci. 121: 747–752.

Neumann M., and M. Affolter, 2006 Remodelling epithelial tubes through cell rearrangements: from cells to molecules. EMBO Rep. 7: 36–40.

Noon E. P.-B., G. Sabarís, D. M. Ortiz, J. Sager, A. Liebowitz, et al., 2018 Comprehensive Analysis of a cis-Regulatory Region Reveals Pleiotropy in Enhancer Function. Cell Rep. 22: 3021–3031.

Nüsslein-Volhard C., and E. Wieschaus, 1980 Mutations affecting segment number and polarity in Drosophila. Nature 287: 795–801.

Okada T., 1954 Comparative morphology of the Drosophilid flies. Phallic organs of the melanogaster group. Kontyu 22: 36–46.

Olesnicky E. C., A. E. Brent, L. Tonnes, M. Walker, M. A. Pultz, et al., 2006 A caudal mRNA gradient controls posterior development in the wasp Nasonia. Development 133: 3973–3982.

Peluffo A. E., I. Nuez, V. Debat, R. Savisaar, D. L. Stern, et al., 2015 A Major Locus Controls a Genital Shape Difference Involved in Reproductive Isolation Between Drosophila yakuba and Drosophila santomea. G3 5: 2893–2901.

Pfreundt U., D. P. James, S. Tweedie, D. Wilson, S. A. Teichmann, et al., 2010 FlyTF: improved annotation and enhanced functionality of the Drosophila transcription factor database. Nucleic Acids Res. 38: D443–7.

Pisarev A., E. Poustelnikova, M. Samsonova, and J. Reinitz, 2009 FlyEx, the quantitative atlas on segmentation gene expression at cellular resolution. Nucleic Acids Res. 37: D560–6.

Polak M., and A. Rashed, 2010 Microscale laser surgery reveals adaptive function of male intromittent genitalia. Proc. Biol. Sci. 277: 1371–1376.

Quinlan A. R., and I. M. Hall, 2010 BEDTools: a flexible suite of utilities for comparing genomic features. Bioinformatics 26: 841–842.

Rafiqi A. M., S. Lemke, S. Ferguson, M. Stauber, and U. Schmidt-Ott, 2008 Evolutionary origin of the amnioserosa in cyclorrhaphan flies correlates with spatial and temporal expression changes of zen. Proc. Natl. Acad. Sci. U. S. A. 105: 234–239.

Rebeiz M., N. H. Patel, and V. F. Hinman, 2015 Unraveling the Tangled Skein: The Evolution of Transcriptional Regulatory Networks in Development. Annu. Rev. Genomics Hum. Genet. 16: 103–131.

Rice G., O. Barmina, D. Luecke, K. Hu, M. Arbeitman, et al., 2019a Modular tissue-specific regulation of doublesex underpins sexually dimorphic development in Drosophila. bioRxiv 585158.

Rice G., J. David, Y. Kamimura, J. Masly, A. Mcgregor, et al., 2019b A Standardized Nomenclature and Atlas of the Male Terminalia of Drosophila melanogaster. Preprints.

Rivera-Pomar R., X. Lu, N. Perrimon, H. Taubert, and H. Jäckle, 1995 Activation of posterior gap gene expression in the Drosophila blastoderm. Nature 376: 253–256.

Russell S. R., N. Sánchez-Soriano, C. R. Wright, and M. Ashburner, 1996 The Dichaete gene of Drosophila melanogaster encodes a SOX-domain protein required for embryonic segmentation. Development 122: 3669–3676.

Sabarís G., I. Laiker, E. P.-B. Noon, and N. Frankel, 2019 Actors with Multiple Roles: Pleiotropic Enhancers and the Paradigm of Enhancer Modularity. Trends Genet. 0. https://doi.org/10.1016/j.tig.2019.03.006

Simmons L. W., 2014 Sexual selection and genital evolution. Austral Entomology 53: 1–17.

Small S., A. Blair, and M. Levine, 1996 Regulation of two pair-rule stripes by a single enhancer in the Drosophila embryo. Dev. Biol. 175: 314–324.

Smith S. J., M. Rebeiz, and L. Davidson, 2018 From pattern to process: studies at the interface of gene regulatory networks, morphogenesis, and evolution. Curr. Opin. Genet. Dev. 51: 103–110.

Staller M. V., C. C. Fowlkes, M. D. J. Bragdon, Z. Wunderlich, J. Estrada, et al., 2015 A gene expression atlas of a bicoid-depleted Drosophila embryo reveals early canalization of cell fate. Development 142: 587–596.

Takahara B., and K. H. Takahashi, 2015 Genome-Wide Association Study on Male Genital Shape and Size in Drosophila melanogaster. PLoS One 10: e0132846.

Takahashi K. H., M. Ishimori, and H. Iwata, 2018 HSP90 as a global genetic modifier for male genital morphology in Drosophila melanogaster. Evolution 72: 2419–2434.

Tanaka K. M., C. Hopfen, M. R. Herbert, C. Schlötterer, D. L. Stern, et al., 2015 Genetic Architecture and Functional Characterization of Genes Underlying the Rapid Diversification of Male External Genitalia Between Drosophila simulans and Drosophila mauritiana. Genetics 200: 357–369.

Tanaka K. M., Y. Kamimura, and A. Takahashi, 2018 Mechanical incompatibility caused by modifications of multiple male genital structures using genomic introgression in drosophila. Evolution. https://doi.org/10.1111/evo.13592

Tomancak P., B. P. Berman, A. Beaton, R. Weiszmann, E. Kwan, et al., 2007 Global analysis of patterns of gene expression during Drosophila embryogenesis. Genome Biol. 8: R145.

Trapnell C., L. Pachter, and S. L. Salzberg, 2009 TopHat: discovering splice junctions with RNA-Seq. Bioinformatics 25: 1105–1111.

True J. R., J. Liu, L. F. Stam, Z.-B. Zeng, and C. C. Laurie, 1997 Quantitative Genetic Analysis of Divergence in Male Secondary Sexual Traits Between Drosophila simulans and Drosophila mauritiana. Evolution 51: 816–832.

Vincent B. J., M. V. Staller, F. Lopez-Rivera, M. D. J. Bragdon, E. C. G. Pym, et al., 2018 Hunchback is counter-repressed to regulate even-skipped stripe 2 expression in Drosophila embryos. PLoS Genet. 14: e1007644.

Whiteley M., P. D. Noguchi, S. M. Sensabaugh, W. F. Odenwald, and J. A. Kassis, 1992 The Drosophila gene escargot encodes a zinc finger motif found in snail-related genes. Mech. Dev. 36: 117–127.

Wieschaus E., and C. Nüsslein-Volhard, 2016 The Heidelberg Screen for Pattern Mutants of Drosophila: A Personal Account. Annu. Rev. Cell Dev. Biol. 32: 1–46.

Wotton K. R., E. Jiménez-Guri, A. Crombach, H. Janssens, A. Alcaine-Colet, et al., 2015 Quantitative system drift compensates for altered maternal inputs to the gap gene network of the scuttle fly Megaselia abdita. Elife 4. https://doi.org/10.7554/eLife.04785

Wunderlich Z., M. D. Bragdon, K. B. Eckenrode, T. Lydiard-Martin, S. Pearl-Waserman, et al., 2012 Dissecting sources of quantitative gene expression pattern divergence between Drosophila species. Mol. Syst. Biol. 8: 604.

Wunderlich Z., C. C. Fowlkes, K. B. Eckenrode, M. D. J. Bragdon, A. Abiri, et al., 2019 Quantitative Comparison of the Anterior-Posterior Patterning System in the Embryos of Five Drosophila Species. G3. https://doi.org/10.1534/g3.118.200953

Yassin A., and V. Orgogozo, 2013 Coevolution between male and female genitalia in the Drosophila melanogaster species subgroup. PLoS One 8: e57158.

Yassin A., and J. R. David, 2016 Within-species reproductive costs affect the asymmetry of satyrization in Drosophila. J. Evol. Biol. 29: 455–460.

Zeng Z. B., J. Liu, L. F. Stam, C. H. Kao, J. M. Mercer, et al., 2000 Genetic architecture of a morphological shape difference between two Drosophila species. Genetics 154: 299–310.

